# Artificial Sweeteners in US-Marketed Oral Nicotine Pouch Products: Correlation with Nicotine Contents and Effects on Product Preference

**DOI:** 10.1101/2024.01.26.577472

**Authors:** Sairam V. Jabba, Peter Silinski, Alicia Y. Yang, Wenyi Ouyang, Sven E. Jordt

**Author notes:** Corresponding author, Address for Corresponding Author, Sven-Eric Jordt, PhD Department of Anesthesiology, Duke University School of Medicine 3 Genome Ct., Durham, NC 27710-3094.

## Abstract

**Introduction:** Artificial sweeteners are listed as ingredients of oral nicotine pouches (ONPs), a new product category with rapidly growing market share. The exact sweetener contents of ONPs remain unknown. Artificial sweeteners in ONPs may facilitate initiation and encourage consumption behavior.

**Aims and Methods:** Artificial sweetener contents in major US-marketed ONP brands (Zyn, on!, Velo) were determined by Liquid Chromatography-Mass Spectrometry (LC-MS). Sweetener effects during the initiation of ONP consumption were modeled in single- and two-bottle tests, offering mice ONP extracts calibrated to contain nicotine levels similar to saliva of people who use smokeless tobacco. To examine the contribution of sweet taste perception, consumption behavior was compared between wild-type mice and mice deficient in the sweet taste receptor (Tas1r2^−/−^).

**Results:** Acesulfame-K was detected in on!, Zyn and Velo ONPs (∼0.3-0.9 mg/pouch), including products marketed as “Unflavored” or “Flavor ban approved”. In Velo ONPs, sweetened with sucralose (0.6-1.2 mg/pouch), higher nicotine strength products contained higher sucralose levels. Tas1r2^−/−^ mice consumed less ONP extracts than wild-type mice in both sexes. ONP extracts with both higher nicotine and sweetener strengths were tolerated by wild-type mice, but produced stronger aversion in Tas1r2^−/−^ mice.

**Conclusions:** ONPs contain significant amounts of artificial sweeteners, with some brands adding more sweetener to ONPs with higher nicotine strengths. Artificial sweeteners, at levels present in ONPs, increase nicotine consumption. Increasing sweetener contents facilitates consumption of ONPs with higher nicotine strengths. Sweetness is a key determinant of ONP use initiation, likely reducing the aversive sensory effects of nicotine and other ONP constituents.

**Implications:** Artificial sweeteners such as acesulfame-K or sucralose reduce aversion and facilitate initiation and continued consumption of ONPs. The marketing of some artificially sweetened ONPs as “Unflavored” of “Flavor ban-approved” suggests that the tobacco industry rejects sweet taste as a determinant for the presence of a characterizing flavor. Sweetness as imparted by artificial sweeteners in tobacco products needs to be addressed by regulators as a component of a characterizing flavor, with the aim to reduce product appeal and initiation by never users, and especially youth attracted to sweet flavors.

## Introduction

Oral nicotine pouches (ONP) are a new smokeless tobacco (SLT) product category with rapidly growing sales in the United States and other countries ^1^. Older SLT product categories such as moist snuff and snus are often sweetened with artificial high-intensity sweeteners such as saccharin or sucralose that are 200-700 times sweeter than table sugar ^2,3^. Some SLT products contain quantities of artificial sweeteners that impart a sweetness more intense than the weight of the product in table sugar ^2^. Sweeteners are also declared as ingredients of ONP, however, the exact sweetener contents in ONP remain unknown ^4^.

Sweeteners have potent behavioral effects, acting through sweet taste receptors expressed in the taste buds of the tongue ^5–7^. Mammalian sweet taste receptors are dimeric G-protein-coupled receptors, consisting of TAS1R2 and TAS1R3 (Taste receptor type 1 subunit 2 & 3) subunits ^5,7^. Genetic deletion of any of the two subunits renders mice incapable of perceiving sweet taste ^5,7^. Sweetened foods and sweet-associated flavors such as vanilla, cotton candy or fruit are strongly preferred by children and adolescents, a preference that extends to flavored tobacco products ^4,8–11^. Tobacco industry documents reveal that tobacco companies carefully adjusted sweetener contents in smokeless products to appeal to users and routinely monitored the sweetener contents of competitors’ products ^12,13^.

Sweeteners such as sucrose (table sugar) or saccharin are routinely used in rodent behavioral studies to increase the oral intake of aqueous nicotine solutions that otherwise would be aversive to rodents ^14,15^. Nicotine, at levels estimated to be present in the saliva of people who use smokeless tobacco products, is perceived as irritating and bitter ^16–20^. The addition of sweeteners is also a widely used strategy to mask irritating and bitter flavors in foods or medicines ^21–23^.

It is unclear whether sweeteners added to ONPs have similar masking effects, suppressing the adverse sensations elicited by nicotine and other ONP constituents released onto the mucosa and into the oral cavity. Such a masking effect can promote product use initiation and consumption by increasing product appeal.

In the present study, we determined artificial sweetener contents of ONPs of the US-market leading brands Zyn, on! and Velo, with different flavors and nicotine strengths. The analysis also included products advertised as “unflavored” or “flavor ban approved” ^24,25^. The role of sweeteners in ONP consumption behavior was examined by comparing the consumption of commercial ONP extracts between wild-type mice and mice deficient in the sweet taste receptor (Tas1r2^−/−^).

## Materials and Methods

### Quantitative analysis of artificial sweeteners in ONPs

Flavored ONP products of the brands Zyn (Swedish Match/Phillip Morris International (PMI)); 9 flavors-“Menthol”, “Cool Mint”, “Peppermint”, “Wintergreen”, “Spearmint”, “Coffee”, “Cinnamon”, “Chill”, and “Smooth”; 3 and 6 mg nicotine strength), on! (Helix Innovations/Altria; 3 flavors; “Mint”, “Cinnamon”, and “Wintergreen” 2 and 8 mg nicotine strength) and Velo (RJ Reynolds/British American Tobacco (BAT); 2 original Velo flavors: Mint and Citrus; 2 and 4 mg nicotine strength; 1 Velo Dryft flavor: Citrus Burst, 4 and 7 mg nicotine strength) (Supplementary Table 1) were purchased from 3 gas stations in Durham, NC between Jan 2020 – Oct 2022. ONP product extracts were prepared by stirring the contents of one pouch overnight at room temperature in 10 mL of water. Extracts were centrifuged and supernatant was collected for chemical analysis of sweeteners. Contents of two high-intensity synthetic sweeteners, acesulfame-k (Sigma Aldrich LLC., Saint Louis, MO) and sucralose (Sigma), declared by the manufacturers as product ingredients, were determined by a modified LC-MS methodology used previously to analyze sweetener contents in snuff and snus, e-cigarettes and other tobacco products (Supplementary methods of Miao et al., 2016).

### Animals

Wild type C57BL/6J mice, male and female (adults aged 8-16 weeks), were purchased from Jackson Laboratories (Bar Harbor, ME). C57BL/6J-backcrossed Tas1r2^−/−^ (taste receptor, type 1, member 2) breeding pairs were obtained from Dr. Richard Pratley (Sanford Burnham Prebys Medical Discovery Institute, FL) with permission from Dr. Charles Zuker (Columbia University, UCSD). Strain congenicity was verified by genome scan SNP analysis (Jackson Laboratories). All mice were maintained at Duke animal facilities in a temperature and humidity-controlled room with a 12-hour light-dark cycle with unlimited access to food and water. All experimental protocols conducted in this study were approved by the Institutional Animal Care and Use Committees of Duke University.

### ONP Product extracts for mouse behavioral studies

ONP product extracts of Zyn Coffee (3 mg nicotine strength) and Velo Citrus (2 and 4 mg nicotine strength) were freshly prepared for drinking experiments on the first day of exposure by stirring the contents of each pouch unit in 20 mL water for 4 hours at room temperature and centrifuged. The supernatants were collected and provided to mice for the drinking studies. Nicotine concentrations in the extracts were estimated at 150 µg/mL for Zyn Coffee (3 mg nicotine in 20 mL water) and either 100 or 200 µg/mL for Velo Citrus (2 or 4 mg nicotine in 20 mL water). These concentrations are predicted to be present in the oral cavity of smokeless product users (Fan et al.,) and determined to produce aversion in mice. Sucralose solutions (0.2%; Sigma) were freshly prepared for each experiment by dissolving sucralose in water. Either amber colored drinking tubes or aluminum foil covered drinking tubes were used to protect nicotine containing ONP extracts from light.

### Mouse drinking assays

Drinking assays with single-housed mice were performed in behavioral chambers (Med Associates, Fairfax, VT) or standard mouse cages with slots for drinking tubes with steel sipper spouts. For acclimation, naïve mice were placed into the chambers overnight (17:00 to 9:00) for 2-3 nights with water only. After acclimation, mice were provided drinking solutions (ONP extracts or nicotine solutions with or without sweetener, or water) on 4 subsequent nights. During acclimation and testing, mice were returned to their home cages during the day, with access to water *ad libitum*. The drinking tubes in each cage were equidistant from food. Consumption volumes were measured by comparing weights of the tubes before and after the overnight drinking periods. Single bottle assay: The consumption of ONP extracts was compared between wild-type and sweet-taste deficient Tas1r2^−/−^ mice using a single bottle assay, with access to extract offered overnight according to the schedule above. ONP extract consumption volumes were normalized to the respective strain controls receiving water only.

Two bottle choice assay: The effects of higher sweetener contents in higher nicotine strength ONP products of certain brands were examined in the two-bottle choice assay and performed according to the schedule above, as described ^20^. Briefly, mice were provided overnight with choice solutions of 2 mg and 4 mg nicotine strength Velo Citrus ONP extracts.

### Statistical analysis

ONP extract consumption volumes were averaged across the 4 overnight sessions. For the single-bottle drinking assay, ONP consumption volumes were normalized to the average volumes of water consumed by the control cohort that were offered plain water only. The unpaired t-test was used to compare ONP extract consumptions between wild-type and Tas1r2^−/−^ mice in the single bottle assays. For the two bottle choice assays, ordinary one-way ANOVA was used to compare daily liquid consumption and preference among groups and Tukey post-hoc was used to compare between individual groups. Further, the paired-t test was used to compare consumed liquid volumes between the two choices within each group. (*p<0.05; **p<0.01; ***p<0.0005; ****p<0.0001). Graphpad Prism (version 10; Boston, MA) was utilized for conducting data and statistical analysis.

## Results

### Artificial sweeteners in ONP products

Artificial sweetener content was determined in the three market-leading ONP product lines, Zyn (PMI/Swedish Match), Velo (BAT, Reynolds, with a subset formerly branded as Dryft) and on! (Altria), across different nicotine strengths and representative flavors. Zyn Chill and Zyn Smooth varieties represented products labelled “Unflavored” or “Flavor-ban approved” ^24,25^. The original Velo nicotine pouch products, Velo Mint and Velo Citrus, contained sucralose, with the average amount of sucralose per pouch increasing with nicotine strength (Figure 1; Top. Velo Mint and Citrus products with 2 mg nicotine strength contained 0.65±0.03 mg/pouch and 0.64±0.03 mg/pouch of sucralose, respectively (N=11 each), while the corresponding products with higher nicotine strength (4 mg) contained approximately twice as much sucralose (“Mint”-1.20±0.06 mg/pouch; “Citrus”-1.19±0.04 mg/pouch; N=11) (Figure 1; Top). One of the Velo Dryft flavors tested, “Citrus burst”, contained acesulfame-k (4 mg: 0.319±0.02 µg/pouch; 7 mg: 0.339±0.02 µg/pouch; N=8) (Figure 1; Top). All the tested Zyn ONP products contained acesulfame-k ranging from ∼0.4 mg - 0.9 mg/pouch (N=4-13), including Zyn Chill and Zyn Smooth, the products labelled “Unflavored” or “Flavor ban-approved” (Figure 1; Bottom). Unlike the original Velo products, sweetener contents in the Zyn products did not increase with nicotine strength. Similar to Zyn ONPs, the on! products contained acesulfame-k, ∼0.65 – 0.94 mg per pouch (N=4-11) (Figure 1; Bottom).

**Figure 1:**
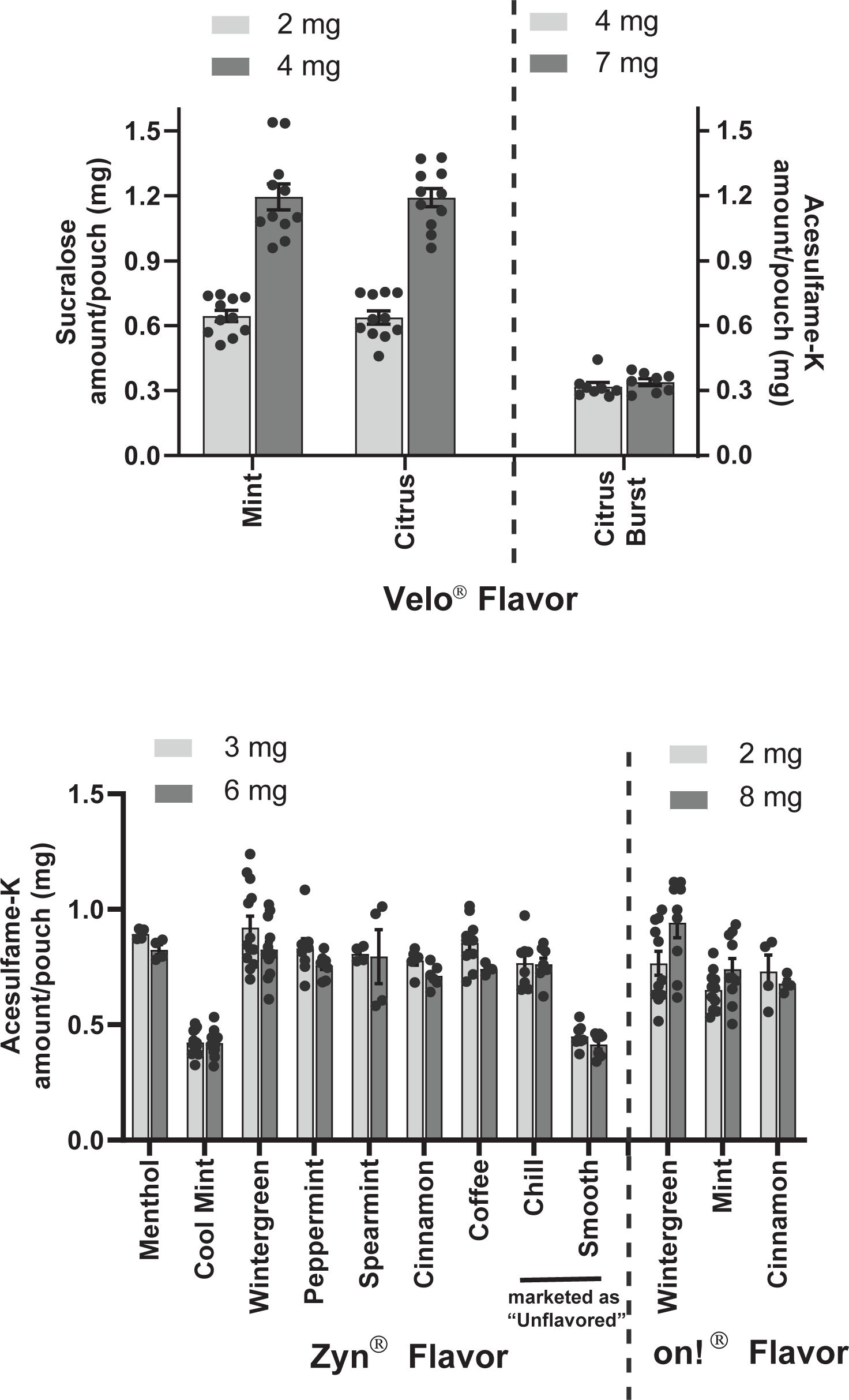
Synthetic sweetener contents in ONP products. Sucralose contents in the two original Velo ONP flavors (RJ Reynolds), Mint and Citrus, for the two available nicotine strengths (2 and 4 mg) and acesulfame-k contents for one of the Velo Dryft flavor, Citrus Burst (4 and 7 mg) (Top panel). Acesulfame-k amounts in Zyn (Menthol, Cool mint, Wintergreen, Peppermint, Spearmint, Cinnamon, Coffee, Chill and Smooth; 3 and 6 mg nicotine strengths; Swedish Match/PMI) and on! (Wintergreen, Mint and Cinnamon; 2 and 8 mg; Altria) ONP products (bottom panel). Sweetener content is represented as mg per unit pouch. Error bars represent SEM. *p<0.05. N=4-13 each.

### Diminished consumption of ONP extracts by sweet taste receptor deficient mice

The effects of sweeteners and sweet perception on ONP extract intake were examined in the single bottle test, comparing consumption volumes of adult female and male wild-type and mice deficient in the sweet taste receptor (Tas1r2^−/−^). In prior control experiments, Tas1r2^−/−^ mice were unable to discriminate between unsweetened and intensely sweetened nicotine solutions that were strongly preferred by wild-type mice (Supplementary Figure 1).

Mice were offered extract from Zyn Coffee-flavored pouches (3 mg nicotine strength) overnight as their only hydration source. The extract was diluted to a nicotine concentration of ∼150 µg/ml, the average concentration estimated to be present in the saliva of smokeless tobacco product users and known to be aversive to mice in both sexes ^20^. Overnight ONP extract consumption volumes were compared between naïve adult wild-type and Tas1r2^−/−^ mice. ONP consumption volumes were normalized to plain water consumption volumes of the respective water control groups (Figure 2). While ONP extract consumption volumes of wild-type mice were similar to their respective water controls (males: 98.8%, N=6-9; females 89.2%, N=6), Tas1r2^−/−^ mice consumed significantly less ONP extract (males: 75.5%, N=6-9; females 70.4%, N=6; p<0.05) (Figure 2).

**Figure 2:**
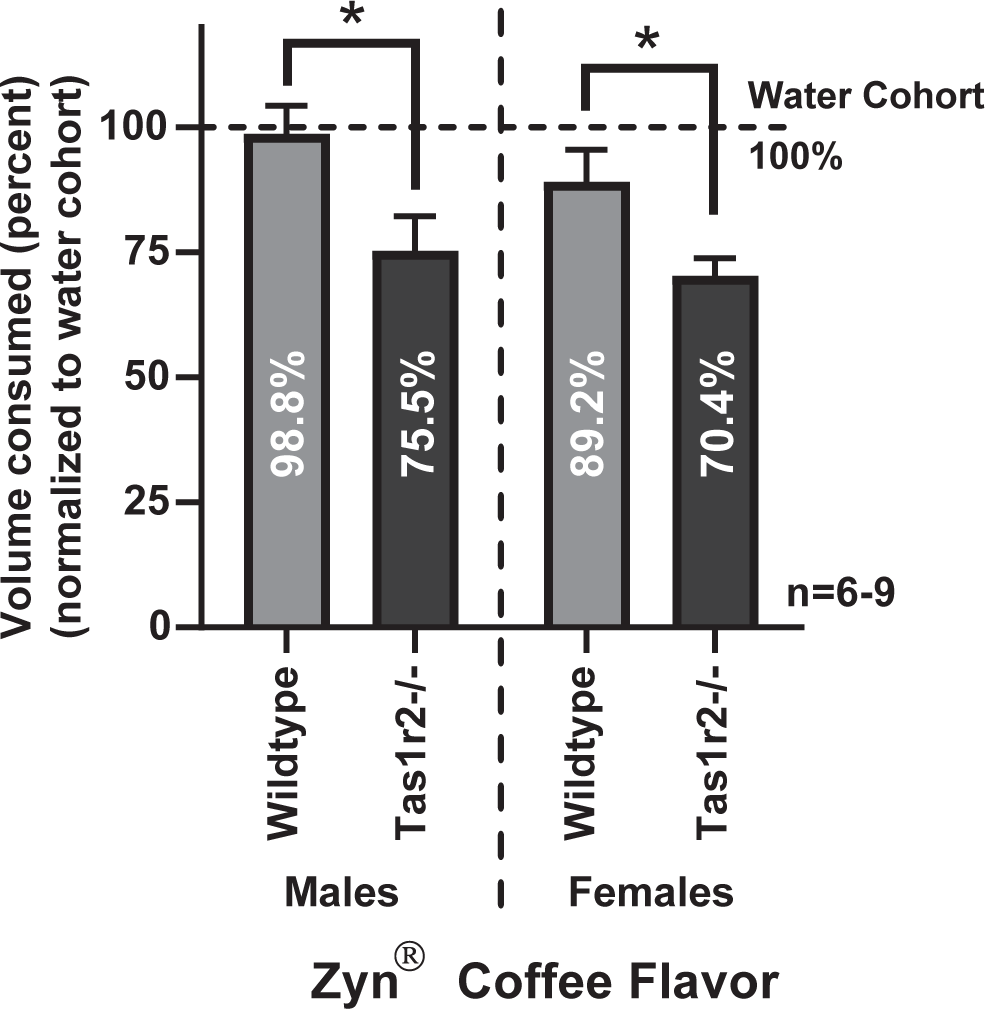
Oral consumption of ONP extract by wild-type mice and mice deficient in the sweet taste receptor, determined in the single bottle test. Normalized consumption volumes of adult male and female C57BL/6J wild-type (light gray) and Tas1r2^−/−^ (black) mice provided with Zyn Coffee (3 mg) extract (nicotine concentration of ∼150 µg/mL), averaged over four nightly testing periods. Data for each strain and sex were normalized to the consumption volumes of mice drinking water only. Normalized percentages are also represented as text inserts in each bar. Bars represent mean±SEM. *p<0.05; N=5-8 per group.

### Facilitation of higher nicotine strength ONP consumption by increased sweetener content

In some of the analyzed ONP product lines we found sweetener contents to be increased in products with higher nicotine strengths (Figure 1; Top). To test whether the increased sweetener contents allow for tolerance of higher nicotine levels, we used the two-bottle-choice test to compare the consumption of extracts of two such products from the Velo Citrus line, with 2 mg and 4 mg nicotine strengths, in wild-type and Tas1r2^−/−^ mice of both sexes. Extracts were diluted 20-fold, resulting in concentrations of ∼100 µg/mL nicotine and ∼30 µg/mL sucralose for Velo Citrus 2 mg, and ∼200 µg/mL nicotine and ∼60 µg/mL sucralose for Velo Citrus 4 mg (Figure 3). Wild-type mice, both female and male, consumed similar amounts of both extracts (Figure 3). In contrast, Tas1r2^−/−^ mice of both sexes consumed larger volumes of the lower nicotine strength (2 mg) product extracts, and smaller volumes of the 4 mg nicotine strength product (Figure 3).

**Figure 3:**
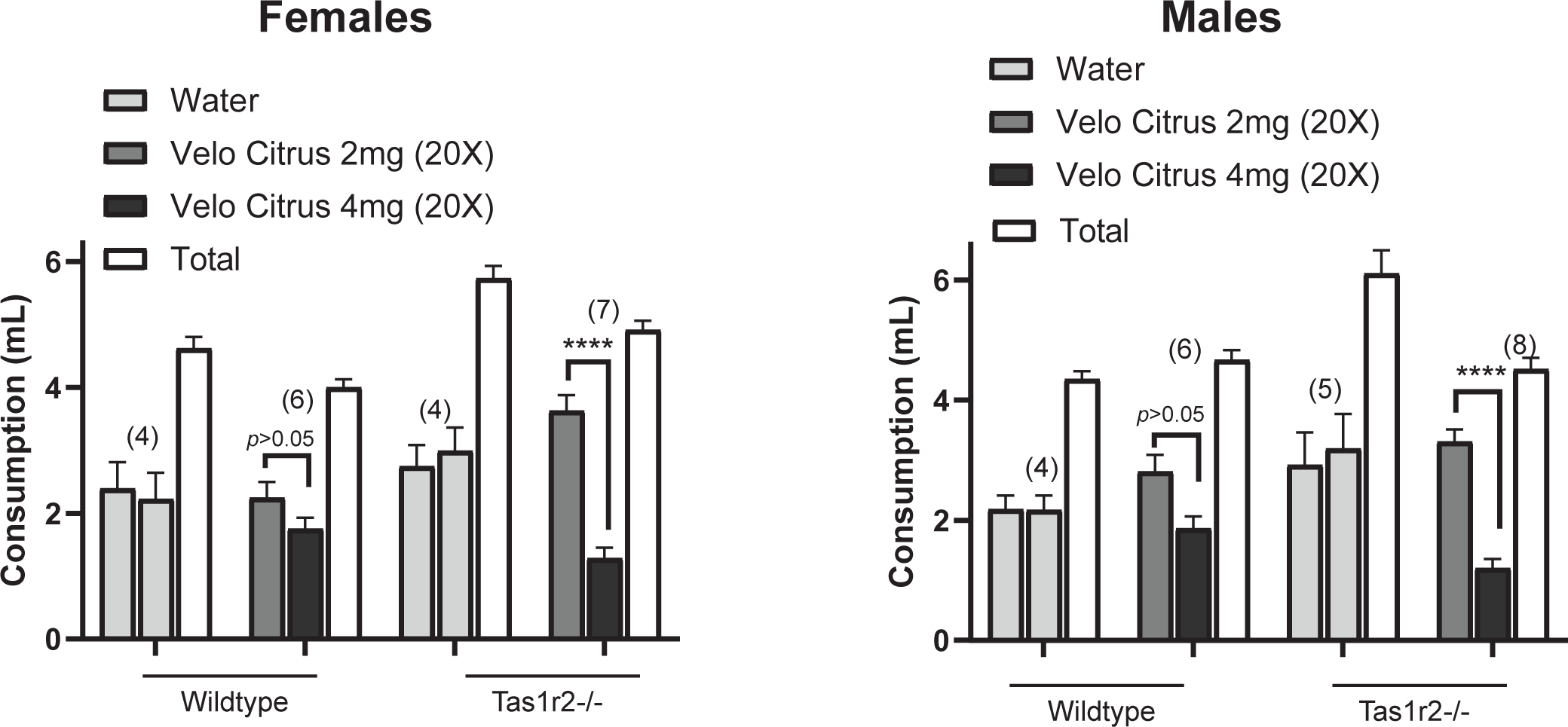
Impact of increased sucralose in higher nicotine strength ONPs on consumption of ONP extracts. Consumption volumes of wild-type and Tas1r2^−/−^ mice in the two-bottle test with choice between extracts of Velo Citrus with 2 mg (dark gray bars) or 4 mg nicotine strengths ((black bars) in females (left panel) and males (right panel). Consumption volumes of control mice presented with plain water only in both tubes are represented in light gray. White bars represent total consumption from both tubes. Consumption volumes were averaged over four nights. Error bars represent SEM. ****p<0.0001. For each group N=4-8/group/sex.

## Discussion

Our chemical analysis revealed that all tested ONP, of all the major US-marketed brands, contained artificial sweeteners. Both Zyn (PMI/Swedish Match) and on! (Altria/Helix Innovations)-branded ONP products exclusively contained acesulfame-k. Velo products originally marketed by RJ Reynolds/BAT contained sucralose. In contrast, the Dryft-branded product line acquired by RJ Reynolds/BAT in 2020 and incorporated into the Velo brand contained acesulfame-k, suggesting that RJ Reynolds/BAT continued manufacturing these separate product lines maintaining their original compositions after the acquisition ^26^.

How does ONP sweetener content compare to other smokeless tobacco product categories? In a previous study, we determined the artificial sweetener contents of snus products marketed by the major tobacco companies (R.J. Reynolds, Altria) in the United States ^2^. We found snus products to be intensely sweetened with sucralose, by far exceeding levels in confectionary products (>6 mg per pouch). In contrast, sweetener contents measured by us in ONP were significantly lower (<1.2 mg per pouch). Snus products contain processed tobacco leaf material, imparting a pungent and acrid flavor due to curing products and alkaloid content. ONP products do not contain tobacco leaf material, with purified nicotine, flavorants and other additives added to a flavor-neutral matrix material. Additionally, snus pouches are heavier and bulkier than ONP pouches, possibly leading to unfavorable release profiles for additives such as sweeteners. For these reasons, it is possible that more sweetener is required to improve the flavor of snus products than ONP.

The mouse behavioral experiments in our study indicate that added sweeteners are strong determinants of product consumption, preference and initiation. When we offered wild-type mice ONP extracts diluted to contain nicotine at the average concentration estimated to be present in the saliva of smokeless tobacco product users (150 µg/ml) ^2^, the animals consumed approximately as much extract as water overnight. This suggests that the flavorants and other constituents of ONPs are calibrated to either mask or tolerate the aversive effects of nicotine at this concentration, known to be strongly aversive when offered in plain water. Mice incapable of tasting sweetness due to genetic deletion of the sweet taste receptor gene (Tas1r2^−/−^) consumed less of the same ONP extracts, suggesting that the sweetener has been added to balance the aversive effects of nicotine and other irritating or bitter constituents.

Our chemical analysis suggests that some manufacturers add higher amounts of artificial sweeteners to ONP products with higher nicotine strengths. While wild-type mice consumed similar volumes of extracts from ONPs with low and high nicotine and sweetener contents, Tas1r2^−/−^ mice consumed significantly less of the extract of the ONP with high nicotine and sweetener contents. These observations support the hypothesis that higher sweetener contents are necessary to effectively mask the aversive effects of the increased nicotine strength.

Since introduction to the US market in 2016, the ONP category has experienced dramatic sales growth ^1,27^. While mint-flavored ONP are the most popular flavor variety, sales of fruit-flavored ONP have increased disproportionately in recent years ^27^. Fruit flavored ONP products are especially popular among youth and young adults who also have a much higher preference for sweetened products than adults ^9,28^. The intense sweetness of artificial sweeteners in ONP, together with added fruit flavors, may increase the risk of initiation of ONP use by youth and young adults who never used tobacco products before. ONP products such as Zyn are virally marketed by “Zynfluencers” on TikTok, YouTube and Instagram, specifically targeting youth and young adults ^29^. These marketing strategies and sales developments resemble those for Juul electronic cigarettes in the years leading up to the youth electronic cigarette epidemic, with sweet fruit flavors the most popular.

In the United States, federal tobacco control legislation has outlawed combustible cigarettes with “characterizing flavors”, with the exception of menthol cigarettes for which a ban has been proposed. The FDA is also regulating the marketing of electronic cigarettes, with marketing approval given to tobacco-flavored products only. Some SLTs have received MRTP (modified risk tobacco product) categorization by FDA, however, the FDA has not made any regulatory determinations on ONPs. The State of California recently banned most tobacco products with characterizing flavors, including ONPs, however, “flavor ban-approved” ONP products continue to be marketed ^24,30^. We detected acesulfame-k in the two Zyn ONP marketed as “flavor ban-approved” or “unflavored”, Zyn Chill and Zyn Smooth, at levels similar to the Zyn products marketed with flavor designators. The fact that these products contain a flavoring agent, the potent artificial sweetener acesulfame-K, is evidence for misleading advertising by the manufacturer. Acesulfame-K is not the only flavorant in these products. In a previous study, we detected a synthetic cooling flavor in Zyn Chill, named WS-3 ^25^, that, while lacking menthol’s minty odor, has similar sensory cooling effects and was previously detected in electronic cigarette liquids and “non-menthol” cigarettes ^31–36^. No cooling agent was detected in Zyn Smooth ONP ^25^.

The fact that the tobacco industry markets artificially sweetened products as “unflavored” or “flavor ban-approved” suggests that the tobacco industry doesn’t consider sweetness conveyed by artificial sweeteners a characterizing flavor as defined by federal and state regulations. This stance contradicts the position of flavor industry organizations such as FEMA, the Flavor and Extract Manufacturers Association, that defines flavor as “… the entire range of sensations that we perceive when we eat a food or drink a beverage. Flavor encompasses a substance’s taste, smell, and any physical traits we perceive in our mouths, …” ^37^. Since sweeteners impart sweet taste, this definition clearly designates sweeteners as flavors. Because of the risk posed by sweetener additives in tobacco products to youth and young adults, legislators and regulators need to determine whether the term “characterizing flavor” in current laws and regulations is defined to encompass sweetness, or whether they need to be amended. Designation of sweetness as a characterizing flavor will allow more effective tobacco product regulation, enabling regulators to limit sweetener contents in tobacco products to minimize risk to public health.

## Acknowledgements

We thank Dr. Ana I. Caceres for support of the behavioral experiments.

## Author contributions were as follows

SVJ and SEJ conceptualized and designed the study; P.S. acquired and analyzed the chemical analytical data; SVJ, WO and AY carried out and analyzed the mouse behavioral experiments; SVJ and SEJ drafted the manuscript; SVJ and SEJ provided supervision.

---

**Supplementary Table 1:** Flavored ONP tobacco products analyzed for synthetic sweeteners in this study, along with their manufacturer, and nicotine strength information.

|  | Brand Name<br>(Manufacturer) | Flavor | Nicotine strength<br>tested (mg/unit ONP) | Synthetic<br>sweetener present |
| --- | --- | --- | --- | --- |
| 1 | Velo (RJ Reynolds) |  |  |  |
|  |  | Mint | 2 and 4 | Sucralose |
|  |  | Citrus | 2 and 4 | Sucralose |
|  |  | Citrus Burst<br>(Velo Dryft) | 4 and 7 | Acesulfame-k |
| 2 | Zyn (Swedish Match/PMI) |  |  |  |
|  |  | Menthol | 3 and 6 | Acesulfame-k |
|  |  | Cool Mint |  |  |
|  |  | Peppermint |  |  |
|  |  | Chill |  |  |
|  |  | Wintergreen |  |  |
|  |  | Spearmint |  |  |
|  |  | Coffee |  |  |
|  |  | Cinnamon |  |  |
|  |  | Smooth |  |  |
| 3 | on! (Altria) |  |  |  |
|  |  | Mint | 2 and 8 | Acesulfame-k |
|  |  | Cinnamon |  |  |
|  |  | Wintergreen |  |  |

**Supplementary Figure 1:**
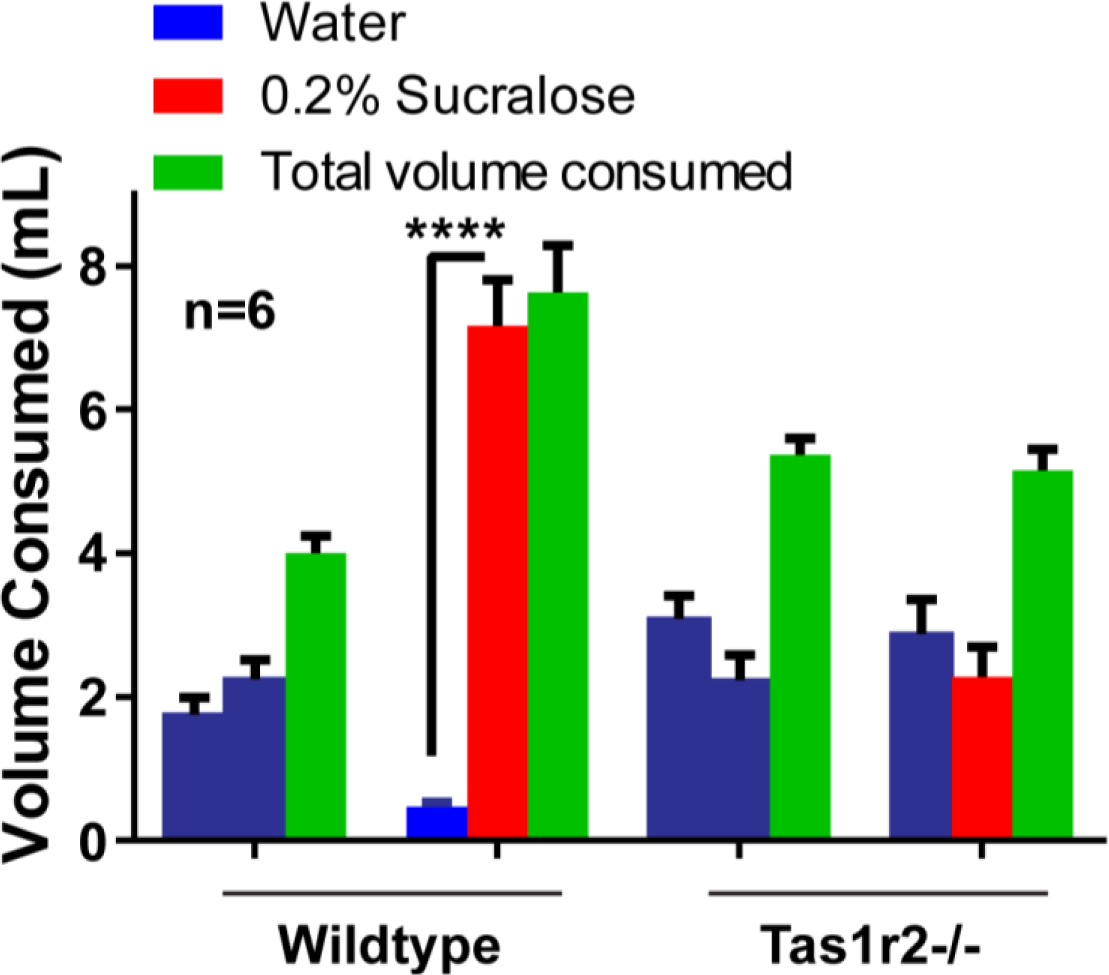
Effect of deficiency of sweet taste receptor Tas1r2 on perception of synthetic sweetener. Consumption volumes of adult C57BL/6 wild-type and Tas1r2^−/−^ mice given the choice between water (blue bars) and sucralose-containing water (2 mg/mL; red), averaged over 4 nightly testing periods. Total consumption from both tubes is also represented (green). Bars represent mean ± SEM. ****:p<0.0001; n=6 per group.

